# Comparative proteomic analysis of the composition of decellularized extracellular matrix (dECM) and dECM-based inks as compared to the native tissue

**DOI:** 10.1101/2024.09.23.614437

**Authors:** Ainhoa Irastorza, Paula Vázquez-Aristizabal, Lore Zumeta-Olaskoaga, Maider Mateo-Abad, Pedro Guerrero, Koro de la Caba, Ander Izeta

## Abstract

Regenerative medicine and tissue engineering approaches based on the use of 3D-bioprinted decellularized extracellular matrix (dECM) present the advantage of a relatively biomolecule-rich matrix, which directs cell growth and differentiation in a tissue-specific manner. However, little is known about the composition changes that occur with standard processing of dECM-based inks. To characterize this process, six porcine tissues/tissue layers (artery, breast, dermis, epidermis, muscle and nerve) were independently decellularized via chemical, mechanical and enzymatic processes and the resulting dECMs formulated into biocompatible inks, to serve as source biomaterials for 3D printing. A comparative liquid chromatography–tandem mass spectrometry (LC–MS/MS)-based proteomic analysis was carried out for native tissue, decellularized and formulated ECMs, and the resulting complexity of the matrisome analyzed. A core matrisome was found to overlap in all decellularized tissues, as well as tissue-specific components that correlated with predicted functional (gene ontology-based) definitions. The proportion of collagens (mostly the α1 chains of collagen type I and III) increased in the final processing step (inks) as compared to the native ECM and dECM stages. Overall, a median of 55 matrisomal proteins (range 45-126) was detected in the dECM-derived inks. This complexity is far superior in terms of mimicking the composition of native tissue to non-dECM-based inks. Our results support the use of dECM-based inks and biomaterials in mimicking native tissue ECM complexity, demonstrating tissue-specific composition, which can improve future therapeutic approximations.

## Introduction

The extracellular matrix (ECM) is a dynamic network of macromolecules arranged according to cell and tissue specificity ^1^. Tissue-resident cells secrete ECM in response to multiple inputs and create a microenvironment of crosslinked proteins and associated regulators, which is collectively known as the matrisome ^2,3^. The matrisome provides an optimal spatial, biophysical and biochemical microenvironment for cell function, which is dynamically regulated by tissue homeostasis ^4,5^. Cells and matrisome thus maintain a bidirectional crosstalk that modulates cell adhesion, migration, proliferation, and differentiation ^1,3,6^, allowing the tissue to adapt to environmental needs.

The development of tissue-engineered constructs for regenerative medicine purposes is based on the use of biocompatible materials that aim to mimic the intricate architecture, mechanical and biological properties of the original healthy tissue ^7–9^. Further adding to the complexity from a materials engineering perspective, engineered constructs will sometimes have to adapt and resist chronically proinflammatory environments where engraftment is needed ^10^. However, and due to their relatively low price and convenience of use, single-component scaffolds (such as gelatin, collagen, or alginate) that poorly mimic the complexity of native tissues are common alternatives to engineer grafts ^3^. Similarly, tumor-derived ECM such as Matrigel®, which is a commercially available product, is often used as a base for hydrogel development. More recently, due to apparent issues with the specification of cells towards the desired phenotype(s), the focus is shifting to biomaterials based on tissue-specific decellularized extracellular matrix (dECM) ^11–15^.

The dECM-based biomaterials are obtained by exposing tissues to chemical (e.g., detergents, hypo/hypertonic solutions), physical (e.g., temperature, mechanical force) and/or biological (e.g., enzymes) agents ^16^. The aim is to remove cells to prevent potential immunological reactions, while minimizing alterations to the biochemical composition, mechanical structure, and potential bioactivity of the tissues ^17^. Decellularized tissues are solubilized through protease and/or chemical treatments, yielding versatile materials suitable for creating foams, films or suspensions. Of note, the digested soluble materials may be used as inks in 3D bioprinting applications ^18–20^. However, this approach presents several shortcomings: (i) dECM-based inks usually present poor mechanical properties that impede their use in extrusion-based bioprinting ^21^, which may be circumvented by the use of alternative 3D printing strategies, such as volumetric bioprinting ^22^; (ii) the components of the ECM are susceptible to the effect of the decellularization agents they are exposed to, affecting their biological potential ^1^; (iii) relatively little is known on the compositional analysis of dECMs and dECM-based inks, which would be needed to develop a deeper understanding of ECM-derived product bioactivity ^11,23^; and (iv) the loss of specific ECM components that are required for proper bioactivity may imply the need of augmentation strategies to recapitulate bioactive, tissue-specific environments^24^. Occasionally, small organ dimensions may impede the extraction of significant ECM quantities, and thus alternative strategies must be sought ^25^.

In this work, we conducted a comparative liquid chromatography–tandem mass spectrometry (LC– MS/MS) analysis of the proteins present in the original tissue ECMs, dECMs, and digested dECM inks, aiming to evaluate the preservation of ECM components throughout the decellularization and solubilization processes. Moreover, this study reports, for the first time, a thorough examination of the matrisome of the ECM, dECM and inks of six relevant porcine tissues or tissue layers. The study unravels the complexity of the matrisome composition and provides relevant information of tissue-specific proteomic hallmarks that may serve for future reference. Finally, key components that could significantly influence cellular response and bioactivity interpretation have been identified for each tissue.

## Results

### Decellularization of native porcine tissues and digestion to formulate inks for bioprinting

The following swine tissues were collected fresh, <2 h after sacrifice of 2 months old Large White pigs, and cryopreserved: aortic artery, *biceps femoris* muscle, breast, sciatic nerve and skin (the latter separated into dermal and epidermal layers). Based on tissue composition, distinct blends of biological and chemical decellularization agents were formulated for each tissue or tissue layer, and combined with physical actuators until satisfactory tissue decellularization standards ^26^ were achieved for each tissue (as detailed in Methods; **Table S1** and **Figure S1**; and in previously published work ^27,28^. Decellularized tissues were digested as specified in **Table S2** to facilitate their use in ink formulation. Biological triplicate samples of native tissue ECM, decellularized tissue ECM (pre-digestion), and decellularized tissue ECM (post-digestion) were then subjected to LC–MS/MS-based proteomic analysis (**Figure 1**).

**Figure 1:**
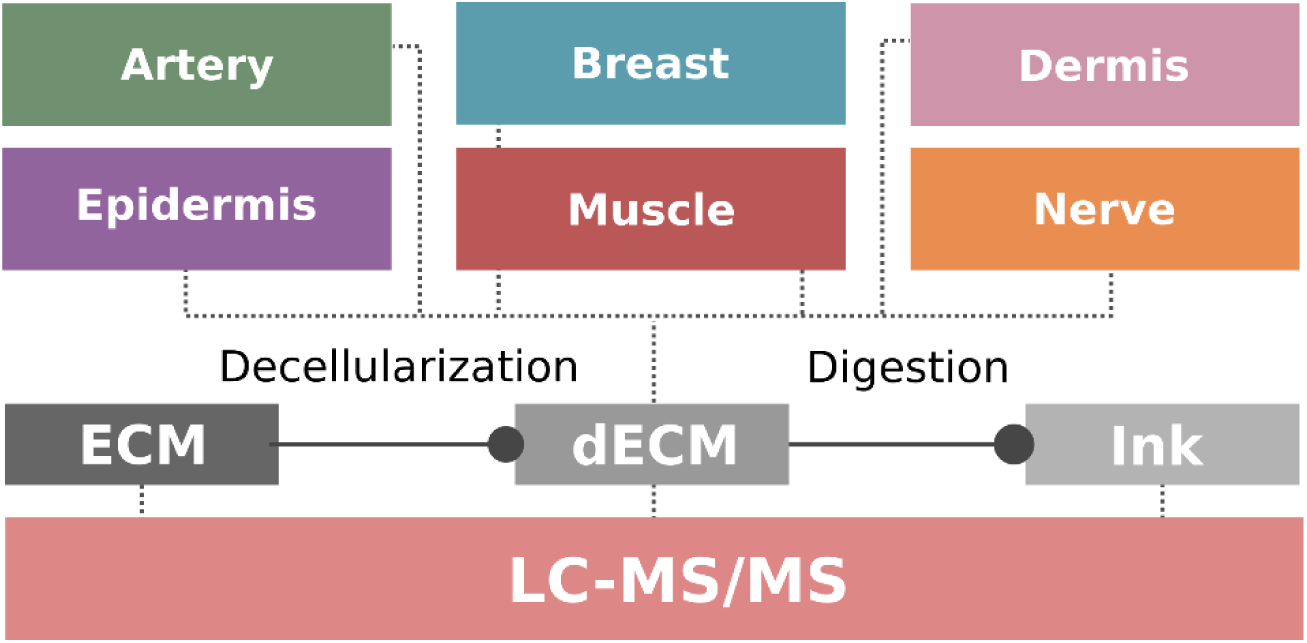
Workflow of the study. Six porcine tissues/tissue layers (artery, breast, dermis, epidermis, muscle, nerve) were independently decellularized and digested, following tissue-optimized protocols. Triplicate samples of the resulting ECM at each step (ECM, dECM, and Ink, respectively) were then analyzed by liquid chromatography–tandem mass spectrometry (LC–MS/MS) to obtain a proteomic atlas of the matrisomal proteins.

### Decellularization and processing induces a distinct loss of ECM-component proteins

To investigate tissue-specific ECM signatures and how well they were conserved after tissue decellularization and sample processing for ink formulation, precursor and fragment tolerances were set at 20 ppm and 0.05 Da, and MaxQuant software searches were carried out against existing databases of pig proteins. Proteins identified in each sample with at least two peptides at FDR < 1 % were further analyzed (peak values are available at PRIDE, identifier number PXD059171). Intracellular and membrane proteins were discarded for further analysis, and ECM-specific proteins were classified by using the MatrisomeDB database ^29,30^, which categorizes ECM proteins into two groups: the “core matrisome” consisting of collagens, proteoglycans, and glycoproteins, and the “matrisome-associated” components (affiliated proteins, regulators and secreted factors), which include ECM-bound proteins and carbohydrates. Matrisomal proteins identified are listed in Zenodo (10.5281/zenodo.14195914).

On average, matrisomal proteins constituted a mere 7.0 % of the total proteins detected in native tissues (160.5 of 2307.0; based on median values of all tissues; **Table S3** and **Figure 2A**). Analysis of dECMs allowed for the detection of 87.0 ± 24.2 % of native proteins, which was perhaps unexpectedly high given the full decellularization obtained, as demonstrated by tissue histology and DNA content analyses (**Figure S1**). In contrast, the same analysis post-digestion (at the ink stage) only detected 23.8 ± 7.3 % of the original tissue proteome (**Table S3**). Of note, the digestion step generates smaller peptides, which may hinder protein identification by LC-MS/MS ^31^. The loss of component proteins with tissue decellularization was clearly tissue-and protocol-dependent (**Figure 2B**). For instance, while arteries, epidermis and muscles retained the total number of identified proteins or even increased it (possibly due to the unmasking of proteins; see below), breast and nerve retained about 70.0 % and the dermis retained only 43.6 % of the total proteins (**Table S3** and **Figure 2B**). The stark difference found in dermal proteome preservation may be due to the use of trypsin at the decellularization step, which was required to eliminate fibroblasts due to the notably ECM-dense nature of the tissue ^32^.

As compared to total proteins, matrisomal proteins seemed to be significantly more resilient to the processing steps, retaining 96.4 ± 19.3 % and 44.1 ± 15.0 % of the native tissue matrisome at the dECM and ink stages, respectively. Decellularization of tissues did not significantly affect detection of matrisomal proteins, as it would be expected due to their extracellular nature. In fact, detection of peptides was increased in several instances (**Table S3**). This could be explained by the fact that many abundant peptides, such as intracellular proteins in native tissue or collagen proteins, mask the intensity of other peptides within the spectrum. As the intensity of the highest peaks decreases, the smaller ones become more apparent ^4^. While every tissue was affected by the digestion step at the matrisome, the epidermis preserved the highest percentage of matrisomal proteins after digestion (72.2 %), which might be due to the different digestion procedure used for this tissue as compared to the other five (see **Table S2**). Within detected matrisome proteins, the most abundant category was glycoproteins, as expected by the large glycoprotein number listed in the core matrisome, followed by the regulators and collagen families (**Figure 2B**) ^33^. As exceptions to this rule, the epidermis and breast tissue-derived ECM, dECMs and specially, inks showed a higher number of matrisomal regulators, followed by glycoproteins and affiliated proteins, which could be possibly explained by the use of tissue-specific protocols that better preserved these biomolecules.

In the light of these findings, Venn diagrams were used to illustrate matrisomal protein composition between ECM, dECM and ink samples for each tissue (**Figure 2C**). The vast majority of the proteins found in the dECM were also present in the ECM, with the exception of breast, dermis and nerve where 18, 46 and 24 proteins were found only in the native samples, respectively. Moreover, some dECMs also revealed uniquely identified proteins; specifically, 10 in epidermis, 6 in muscle, 5 in artery, 3 in breast and 2 in dermis tissue. Likewise, most of the proteins present in the inks were shared among the three groups, although a few unique proteins were also identified.

Finally, it is important to note that dECM-based inks included a median of 53 matrisomal proteins (range 39-122; **Table S3**). This represents a staggering complexity, far superior in terms of mimicking the composition of native tissue to non-dECM-based inks.

**Figure 2:**
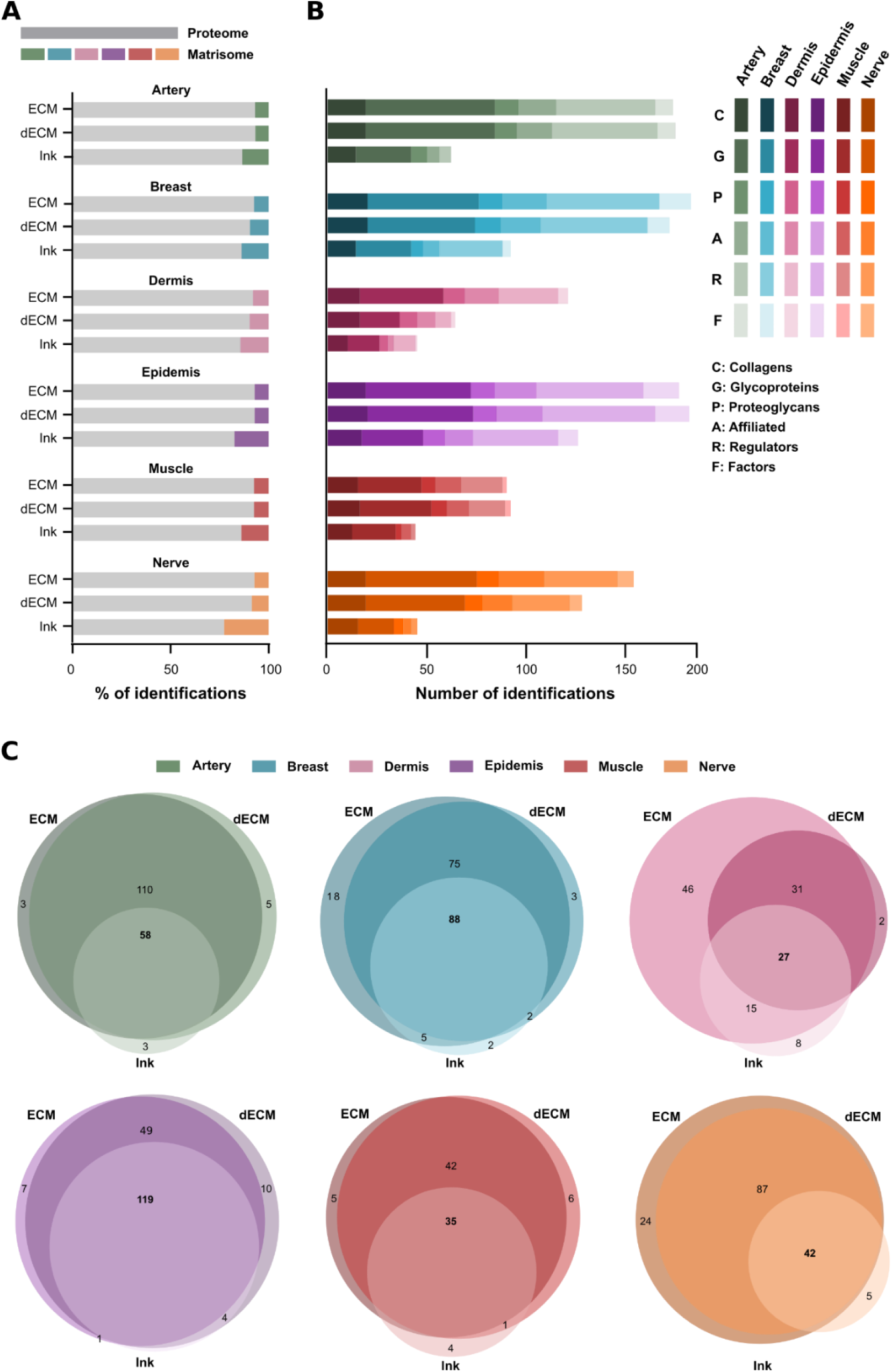
Representation of the contribution as well as conservation of the matrisome and its different categories for the different tissues and samples. **A)** Quantification of the matrisome proportion within the whole proteome of artery (green), breast (blue), dermis (purple), epidermis (violet), muscle (brown) and nerve (orange). **B)** Categorization of the identified proteins by matrisome category, tissue, and step. Distribution of the proportions of MatrisomeDB categories are shown for the artery, breast, dermis, epidermis, muscle and nerve. Top, middle and bottom bars show the patterns for native tissue ECM (ECM), decellularized tissue ECM (dECM) and digested decellularized tissue ECM (Ink), respectively. MatrisomeDB categories are shown in decreasing color intensities. **C)** Venn diagrams of the identified proteins in each case, displaying the overlapping proteins and the proteins unmasked by tissue processing.

### Proteomic signatures of ECMs share a common core of matrisomal proteins

To further analyze the similarities and differences among matrisomal protein profiles detected in each tissue, the list of matrisome identifications was manually supervised to quantify the overlap of shared proteins among tissues (**Figures 3** and **S2**). A Venn diagram shows the overlap of identified proteins found within the ECM of each tissue (**Figure 3A**). A total of 67 proteins were commonly shared among the samples, which in a Gene Ontology (GO) analysis appeared to be related to ECM organization, collagen fibril organization, extracellular structure organization, external encapsulating structure organization, and supramolecular fiber organization (**Figure S3**). These are functions commonly attributed to matrisomal core proteins. Notably, the arterial matrisome presented 32 unique proteins, while other tissues showed comparably lower numbers of unshared proteins. Overall, an overlap of 54.5 ± 11.2 % of matrisome proteins was found among tissues, being lowest in-between artery and muscle (42.1 %), and highest in-between breast and epidermis (87.1 %) (**Figure S2**).

The above results were based on the detection or absence of proteins, and they did not take into account the intensity of the peaks obtained by LC-MS/MS. To account for this variable, a multivariate dimensionality-reduction tool was used to convert multidimensional data to a lower dimensional space. To this end, raw data were subjected to Partial Least-Squares Discriminant Analysis (PLS-DA) (**Figure 3B**) ^34,35^. Together with this, in order to unveil the major proteins contributing to inter-tissue differences, a PLS-DA-based Variable Influence/Importance on Projection (VIP)-scoring analysis was conducted for every component explaining the covariance (**Figure 3C**) ^36,37^. The PLS-DA analysis demonstrated a clear discrimination of tissue-specific signatures, as observed in the distribution of groups based on the principal components. According to the axes defined by components 1 (PC1) and 2 (PC2), there was a clear distinction, on one hand, between epidermis and artery samples, and on the other hand, between these and the rest of the tissues. Based on the results of the VIP score analysis, it was shown that, for both PC1 and PC2, the proteins with the greatest contribution to the observed differences included filaggrin (*FLG*), anti-mullerian hormone (*AMH*), cathepsin H *(CTSH*), galectin 7 (*LGALS7*), secretory leukocyte peptidase inhibitor (*SLPI*) as well as numerous proteins within the S100 (*S100A2, S100A6, S100A9* and *S100A16*), serpin(*SERPINB2, SERPINB5, SERPINB7, SERPINB8, SERPINB10* and *SERPINB13*), and annexin families (*ANXA1, ANXA2, ANXA8* and *ANXA9*). These proteins exhibited higher abundance in the epidermis group and perform structural and functional roles in the epidermal barrier (*FLG2, COL17A1, LAMC2*, and *LAMA3*); differentiation and maintenance of keratinocytes (*SERPINB13, ANXA8* and *TGM5*); stress response and skin repair (*S100, LGALS7, ANXA2, SLPI* and *ANXA1*); as well as remodeling of the ECM (*CTSH* and *CTSA*). Conversely, *FMOD, PRELP* and *GPC6* were overexpressed in the artery, contributing to the inter-tissue differences. In this case, both fibromodulin (*FMOD*) and prolargin (*PRELP*) have roles in ECM structuring. The first specifically regulates the organization of collagen type I and type III fibers, while the second promotes interaction between collagen and other ECM components, such as laminins and fibrillins. Additionally, glypican-6 (*GPC6*) is a membrane proteoglycan that regulates cellular signalling by modulating growth factor pathways such as Wnt. The breast sample demonstrated an overexpression of different proteins grouped within the axis of component 3 (PC3), which enabled its discrimination from the other tissues. Among these proteins were filaggrin 2 (*FLG2*), S100 calcium binding protein A6 (*S100A6*), serpin family B member 13 (*SERPINB13*), and anti-mullerian hormone. Finally, the axis defined by component 4 (PC4) primarily allowed for the distinction between dermis and muscle samples relative to nerve samples, due to an overexpression of proteins such as Von Willebrand Factor A domain containing 1 (*VWA1*), ficolin 2 (*FCN2*), nidogen 2 (*NID2*), S100 calcium binding protein B (*S100B*), α1 chain of collagen XXVIII (*COL28A1*), and ADAM metallopeptidase domain 10 (*ADAM10*) in the last tissue. The proteins that contributed to these differences are involved in various functions such as ECM remodeling and cell adhesion (*VWA1, NID2, COL28A1, FN1, MATN2, LAMA1, HMCN2, EMILIN3* and *COL14A1*), cell signaling and neural development (*ADAM10, GPC4, PLXDC2* and *WNT5B*), damage response and neural protection (*FCN2, S100B, SERPIND1, SERPINA3, KNG1, ITIH4* and *CNTF*), as well as the regulation of the blood-brain barrier and vascular integrity (*LAMA1* and *COL18A1*) ^66,67^.

Thus, PLS-DA analysis revealed that even if the proteins are shared among the ECMs of different tissues, the proportion and relevance of said proteins is different, allowing to define separate groups, probably as a consequence of tissue-specific matrisomes. Overall, we found substantial overlap of core matrisomal proteins in-between native tissues, as well as the existence of a distinct proteomic signature for each tissue.

**Figure 3:**
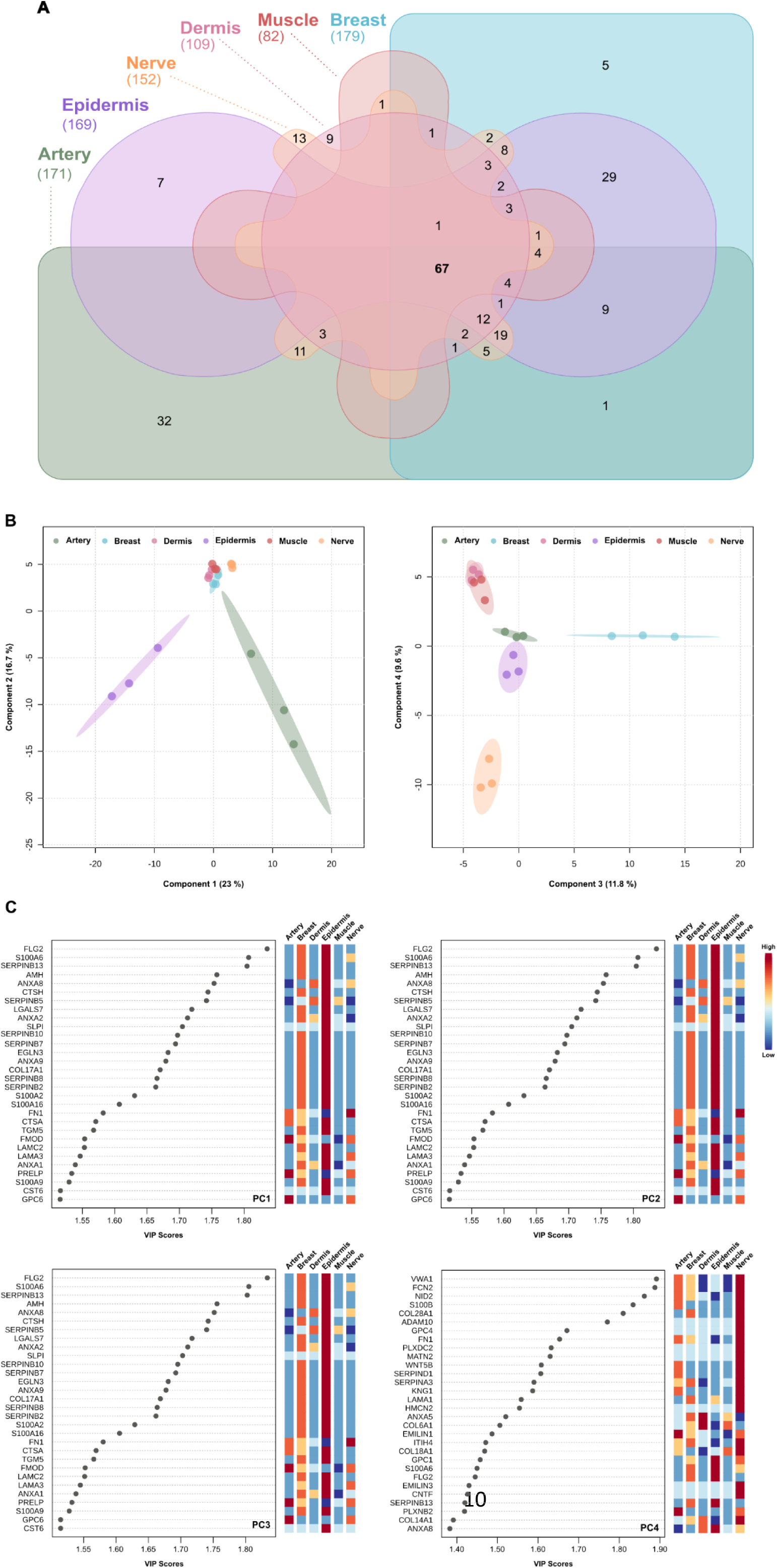
Analysis of the similarities between native-ECM matrisomal protein profiles. **A)** Venn diagram of the distribution of the shared proteins of artery (green), breast (blue), dermis (purple), epidermis (violet), muscle (brown) and nerve (orange). **B)** Partial Least-Squares Discriminant Analysis (PLS-DA) analysis of same tissues. **C)** PLS-DA-based Variable Influence/Importance on Projection (VIP)-score of the major contributors to the differences among tissues.

### Composition of the core matrisomal proteomes of ECMs

With the intent of understanding the composition of the matrix in the native tissues, dECMs and inks, the profile of the matrisomal proteins (as defined by the different categories of MatrisomeDB) was plotted by averaging peak intensity (**Figure 4**). Overall, while all the native tissues showed a great diversity of proteins of the different matrisome categories, this diversity was reduced after the decellularization step, with proteoglycans and matrisome-associated proteins (affiliated, regulators and factors) being particularly affected. This may be due to the fact that this type of proteins are hydrophilic and thus more susceptible to denaturation triggered by decellularizing agents and digestion procedures ^39,40^. Unlike collagens, that usually compose the main structure of tissues, proteoglycans and matrisome-associated proteins play more dynamic roles acting as intermediaries between cells and their environment ^41^. In addition, the triple helix configuration of collagen and the carbohydrate chains present in the glycoproteins, may have provided greater stability to these proteins, allowing a certain enzymatic resistance that contributed to their preservation ^42^. Accordingly, the inks retained mainly collagens and glycoproteins, except for the epidermis.

As expected, there were some differences in-between tissues. Both arterial ECM and dECM showed a heterogeneous protein profile, with diverse presence of all matrisome categories. However, the digestion process induced a reduction of the associated proteins along with the proteoglycans, with a final composition based mainly on collagens (58.9 %) and glycoproteins (32.7 %). The breast tissue ECM showed an equilibrated matrisome, with high abundance of associated matrisome proteins (37.6 %). The core was composed of 28.6 % collagens, 22.7 % glycoproteins and 11.1 % proteoglycans. The amount of core proteins augmented to 92.5 % in the case of dECM, and 92.2 % in the case of the ink. Interestingly, although most detected proteins were related to the core matrisome, a 2.5 % of regulators was present in the breast ink.

**Figure 4:**
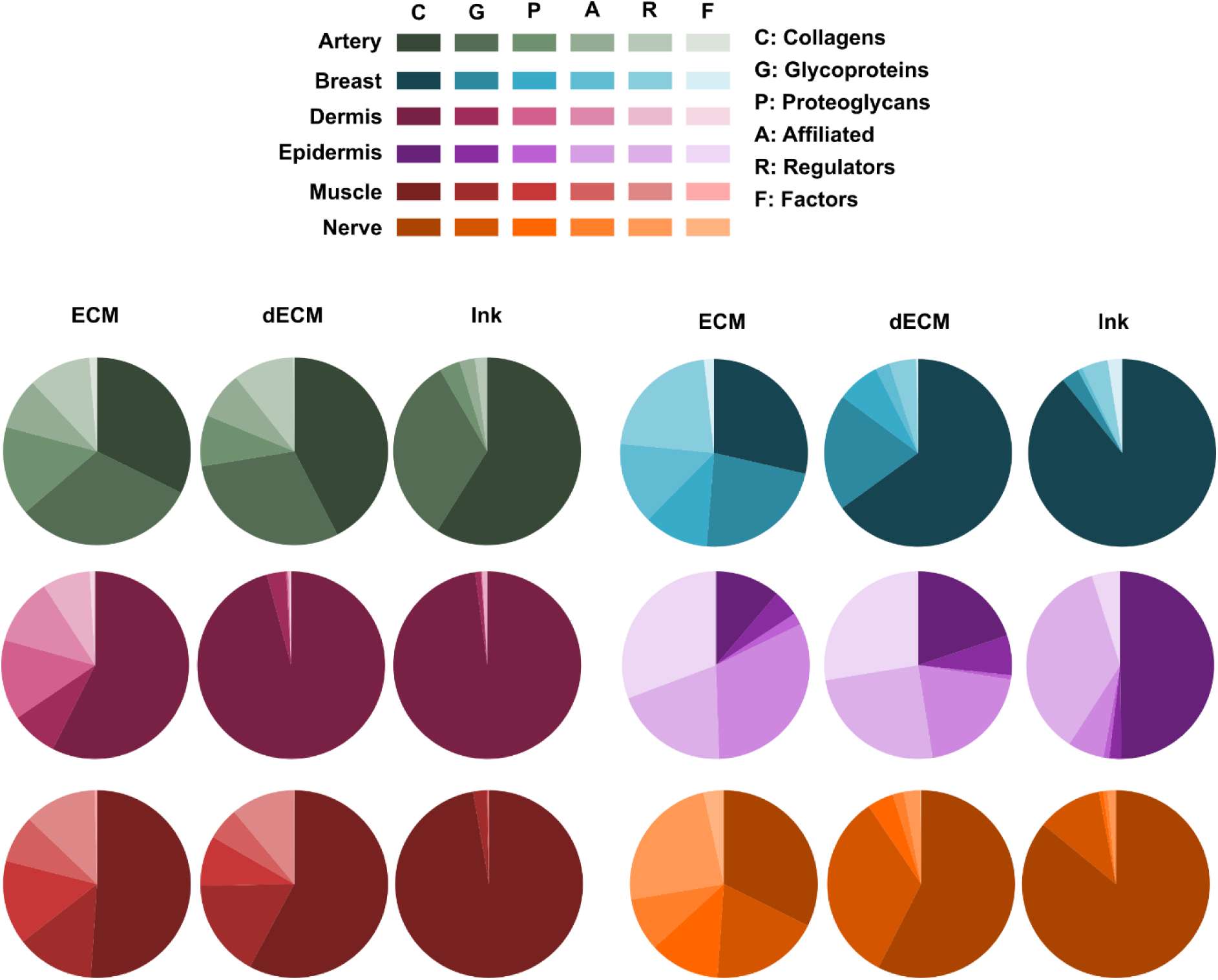
Proportions of the different matrisomal protein categories in each tissue at the ECM, dECM and ink processing stages. Distribution of the proportions of MatrisomeDB categories are shown in pie charts for the artery (green), breast (blue), dermis (purple), epidermis (violet), muscle (brown) and nerve (orange). Left, middle and right columns show the patterns for native tissue ECM (ECM), decellularized tissue ECM (dECM) and digested decellularized tissue ECM (Ink), respectively. MatrisomeDB categories are shown in decreasing color intensities as follows: Matrisome core: C (Collagens), G (Glycoproteins), P (Proteoglycans) and matrisome-associated: A (Affiliated), R (Regulators), and F (Factors).

The main contribution in the dermal ECM was given by collagens (57.4 %), followed by 13.8 % of proteoglycans. Processing induced an increase in collagens that accounted for 95.8 % in dECM and 97.9 % of the total intensity in the Inks. Collagens were followed by the glycoprotein category as second most abundant in both cases. Relatively small changes were observed in the digestion step (dECM to ink), possibly because the major proteome loss could be attributed to the trypsin treatment necessary to release the cells from the dermal ECM at the decellularization step. Due to the high abundance of collagen family proteins, a distribution of the different collagen types per tissue and sample type was calculated. Although collagen type I was predominant in the inks, 16 collagen isoforms were detected, with significant inter-tissue diversity (**Figure S4**).

An exception to collagen predominance was the epidermal ECM (11.2 % collagens). A significant proportion of this ECM consisted of affiliated proteins (31.6 %), regulators (19.8 %), or secreted factors (30.7 %), suggesting a particularly low presence of core matrisome proteins in this tissue. However, epidermal ink composition included 52.9 % core proteins (49.8 % of which corresponded to collagens) and 47.1 % matrisome associated proteins, respectively. Intriguingly, 36.0 % of the epidermal ink were regulator proteins. Even if keratins play a significant role in the epidermal microenvironment, these were not considered for analysis in this study, as they are typically considered as contaminants and removed in the pre-processing of the LC-MS/MS data of every tissue.

Native muscle tissue was mainly composed of collagens (51.0 %), proteoglycans (14.5 %), glycoproteins (13.4 %) and regulators (12.5 %). The heterogeneity and contribution of the different matrisome categories was maintained after decellularization, although the proportion of proteoglycans decreased slightly (8.6 %). More notably, after the digestion process, only a small fraction of the original glycoproteins (2.3 %) was retained, resulting in a composition mainly based on collagens, which accounted for 97.2 % of the identified proteins.

Similar to breast tissue, nerve sample revealed a balanced protein composition composed mainly of collagens (32.3 %), regulators (24.2 %), glycoproteins (18.8 %) and proteoglycans (12.1 %). The decellularization protocol had an impact on proteome diversity by reducing the presence of proteoglycans and matrisome-associated proteins, increasing collagens (57.5 %) and glycoproteins (33.0 %) instead. This phenomenon was more pronounced in nerve-derived inks, where proteoglycan groups, affiliated proteins, regulatory and secreted factors represented only 2.9 % of the proteome composition.

### Native tissue-specific proteomic signatures of ECMs partially align with GO terms

To further delineate tissue-specific proteomic signatures, a heatmap showing upregulated and downregulated clusters per tissue of the top 100 most abundant matrisomal proteins was construed for native tissue ECMs (**Figure 5**). Of interest, arteries, breasts, epidermis, muscle and nerves showed specifically upregulated protein clusters of 8-30 proteins that were consistently found in the biological triplicates. In contrast, the dermis showed a more generalized pattern of protein downregulation, indicating that its ECM composition was comparatively non-specific.

The genes encoding the proteins that were uniquely upregulated in each tissue were subjected to GO analysis. The proteins found clustering for the epidermal ECM signature corresponded to general ECM GO terms such as regulation of peptidase activity, but also included tissue-specific terms such as epidermis development. In the case of the arteries, in addition to common ECM-related GO terms the identified proteins were related to tissue-specific processes such as regulation of plasminogen activation, positive regulation of hemostasis and coagulation. Breast tissue ECM cluster was found to be related to more general terms such as negative regulation of blood coagulation and fibrinolysis. The proteins clustering for neural tissue were related to general ECM-related terms, as well as to the Wnt signaling pathway and notochord development. The ECM signature corresponding to the muscle was related to general ECM terms such as regulation of basement membrane organization or positive regulation of integrin-mediated signaling pathway, but terms related to muscle cell differentiation were also identified. Detailed results are presented in **Table S4**.

In summary, clusters with more complex compositions tended to be associated with more specific GO terms, while clusters characterized by general GO terms were often indicative of less well-defined clusters.

**Figure 5:**
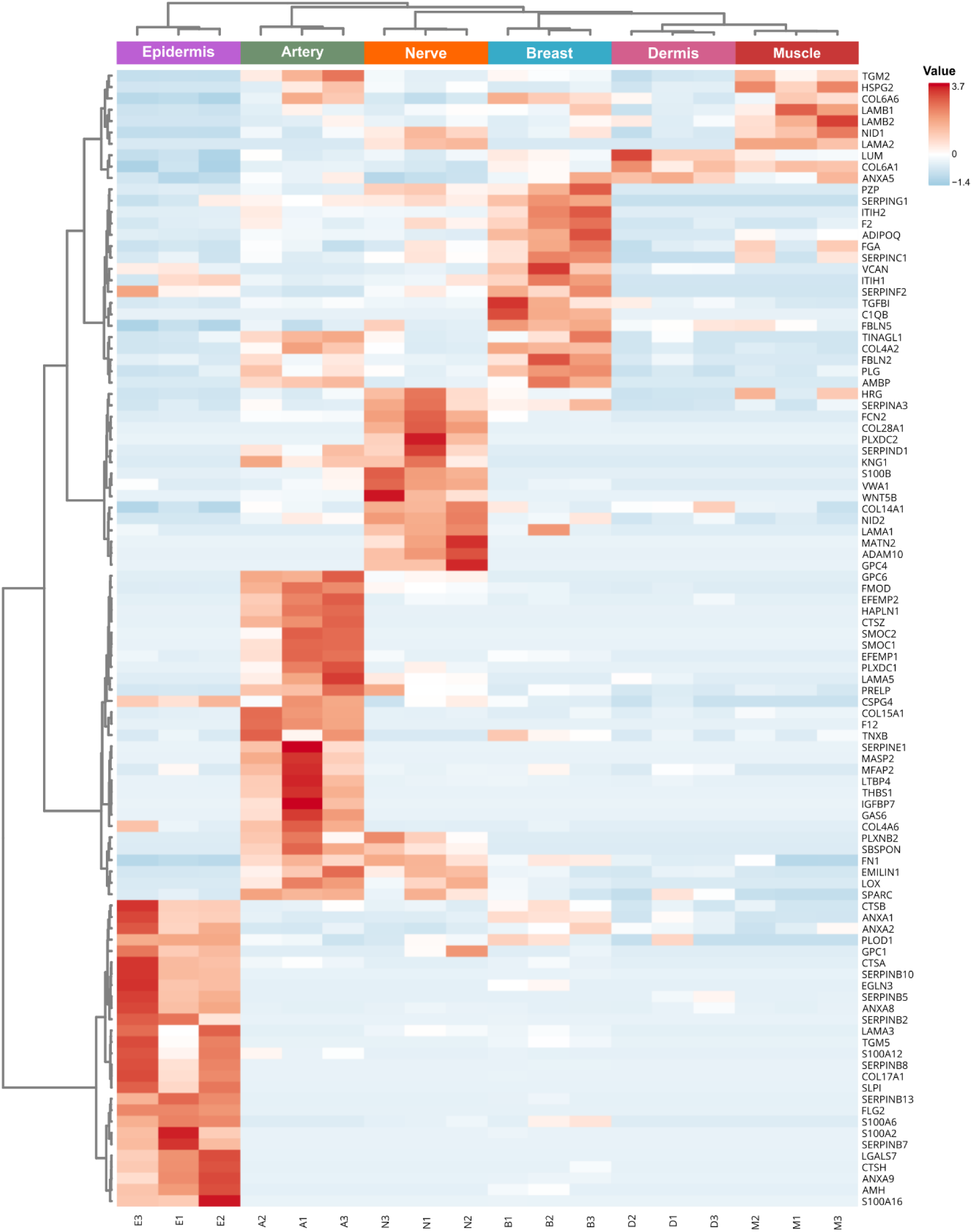
Heatmap of matrisome proteins of each tissue ECM. The top 100 most abundant proteins are shown, clustered according to their contribution to the protein profiles of the different tissues and biological samples. In addition, dendrograms showing the clustering relationships both in-between samples and in-between proteins are plotted.

### Proteomic signatures of ECMs at the differing processing steps scatter in PLS-DA analysis as a function of a limited number of up-and downregulated proteins

To quantitatively capture the inter-group variation in protein composition among the processing steps, a cross-validated PLS-DA and VIP-scoring analysis were conducted using the normalized average label-free quantitation (LFQ) intensity proteomic data from the ECM, dECM, and ink processing stages of each tissue (**Figure 6**). The results of the cross-validation are represented in Figure S6.

In all the studied tissues, the component 1 axis (PC1) distinguished between the ECM sample and the ink sample, while the component 2 axis (PC2) differentiated the decellularized matrix from the other two samples.

For the artery, the proteins contributing to the differences included the α1 and α3 chains of collagen VI (*COL6A3* and *COL6A1*), the α1 chain of collagen XV (*COL15A1*), and transforming growth factor beta induced (*TGFBI*), which were overexpressed in the native sample. In the dECM, there was an increase in the α6 chain of collagen IV (*COL4A6*), as well as proteins such as HtrA serine peptidase 3 (*HTRA3*) and procollagen-lysine,2-oxoglutarate 5-dioxygenase 2 (*PLOD2*). The final sample exhibited a higher abundance of the α1 and α2 chains of collagen I (*COL1A1* and *COL1A2*) together with fibulin 5 (*FBLN5*). Validation revealed some dependency on replicates regarding components 3 and 4, making it hard to reach clear conclusions.

The VIP-scoring analysis demonstrated that in both breast and nerve tissues, all proteins encompassed in PC1 were overexpressed in the ECM, while those in PC2 were overexpressed in the dECM. In both cases, the ink sample exhibited downregulation of the identified proteins. In breast tissue, proteins overexpressed in the ECM included members of the annexin family, specifically A11, A6, A7, and A4 (*ANXA11, ANXA6, ANXA7* and *ANXA4*), serpins F1 and G1 (*SERPINF1* and *SERPING1*), and other proteins such as plasminogen (*PLG*) and decorin (*DCN*). In the decellularized breast sample, there was an increased abundance of collagens, specifically the α1, α2, and α6 chains of collagen VI (*COL6A1, COL6A2* and *COL6A6*), the α2 and α6 chains of collagen IV (*COL4A2* and *COL4A6*), and the α1 chain of collagen XXI (*COL21A1*).

In the nerve ECM sample, overexpressed proteins included annexins A1 and A11 (*ANXA1* and *ANXA11*), serpins G1, H1, and C1 (*SERPING1*, *SERPINH1*, and *SERPINC1*), and nidogens 1 and 2 (*NID1* and *NID2*). In contrast, the dECM exhibited higher levels of fibrillins 2 and 3 (*FBN2* and *FBN3*), the α3 chain of collagen VI (*COL6A3*), the α6 chain of collagen IV (*COL4A6*), and the α1 chains of collagens III, VIII, and XV (*COL3A1, COL8A1* and *COL15A1*). Other overexpressed proteins in the dECM included laminin subunit α5 (*LAMA5*), versican (*VCAN*), and elastin (*ELN*).

Analysis of dermis samples revealed that the proteins contributing to variability along PC1 included the α1 and α2 chains of collagen I (*COL1A1* and *COL1A2*), which were overexpressed in the ink sample. The remaining proteins within this component were more abundant in the native sample, including annexins A5 and A2 (*ANXA5* an*d ANXA2*), fibronectin 1 (*FN1*), fibulin 5 (*FBLN5*), the α1 chains of collagens VI and XIV (*COL6A1* and *COL14A1*), the α6 chain of collagen VI (*COL6A6*), and several laminins (*LAMC1, LAMB2*, and *LAMA4*). Proteins in component 2 showed that the decellularized dermis sample had higher expression of dermatopontin (*DPT*) and the α1 chains of collagens (*COL21A1* and *COL5A1*). Similar to the artery, components 3 and 4 depended on biological replicates, limiting conclusions.

In the epidermis, similar to breast and nerve samples, all PC1 proteins were overexpressed in the native sample. These included transglutaminase 2 (*TGM2*), inter-alpha-trypsin inhibitor heavy chain 2 (*ITIH2*), latent transforming growth factor beta binding protein 2 (*LTBP2*), and families such as serpins (*SERPINB5, SERPINF1, SERPINB10* and *SERPINH1*) and annexins (*ANXA2* and *ANXA8*). Similarly, the dECM showed overexpression of most genes associated with PC2, including proteoglycan 3 (*PRG3*), extracellular matrix protein 1 (*ECM1*), thrombospondin 4 (*THBS4*), transglutaminase 1 (*TGM1*), and cystatin A (*CSTA*).

Lastly, in muscle tissue, the native sample showed overexpression of PC1 proteins, including the α1, α2, α3, and α6 chains of collagen VI (*COL6A1, COL6A2, COL6A3* and *COL6A6*), as well as lumican (*LUM*), heparan sulfate proteoglycan 2 (*HSPG2*), transglutaminase 2 (*TGM2*), and osteoglycin (*OGN*). Conversely, the ink sample had higher levels of the α1 and α2 chains of collagen I (*COL1A1* and *COL1A2*), the α2 chain of collagen IV (*COL4A2*), and the α1 chain of collagen III (*COL3A1*). The decellularized sample exhibited overexpression of most PC2 proteins, such as annexin 8 (*ANXA8*), the α1 chains of collagens IV and VII (*COL4A1* and *COL7A1*), EGF containing fibulin extracellular matrix protein 2 (*EFEMP2*), AE binding protein 1 (*AEBP1*), and microfibril associated protein 2 (*MFAP2*).

**Figure 6:**
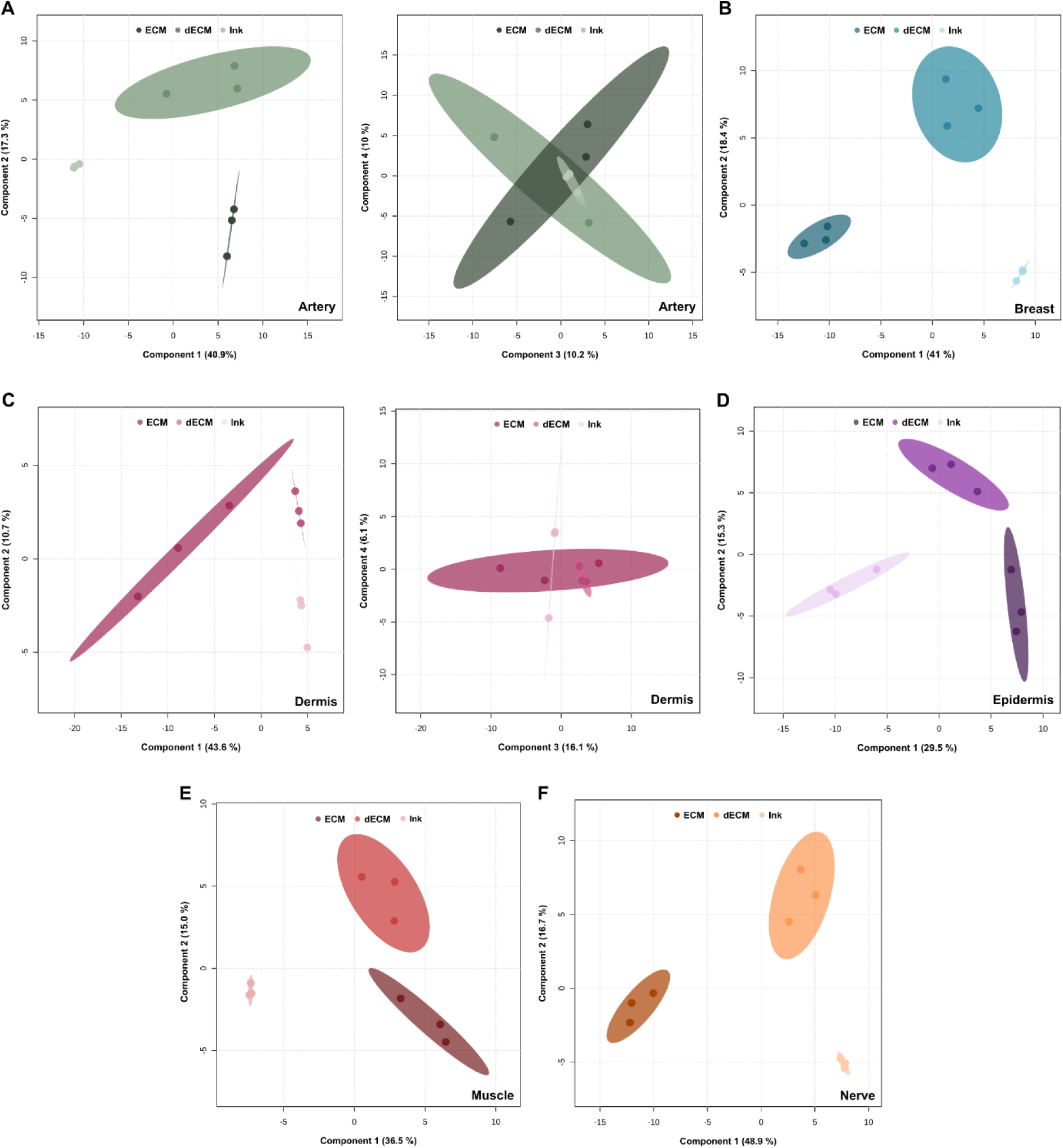
Partial Least-Squares Discriminant Analysis (PLS-DA) analysis of the normalized average label-free quantitation (LFQ) intensity proteomic data from ECMs, dECMs and inks. Distribution of samples are shown in decreasing color intensities for the ECMs (dark colors), dECMs (intermediate colors) and inks (light colors), for the following tissues: **A)** artery (green), **B)** breast (blue), **C)** dermis (purple), **D)** epidermis (violet), **E)** muscle (brown), and **F)** nerve (orange).

### Some tissues are more heterogeneous than others with regard to composition of abundant proteins

To delve further into the exact protein composition of ECMs, we depicted 98 of the proteins that displayed an abundance >1 % in each tissue, aiming to ascertain if there were distinct patterns unique to each tissue type (**Figure 7**). The rest of abundant proteins were commonly classified as “Other”.

As expected, every tissue displayed a unique signature when plotting proteins by abundance. Arteries, breast and nerve-derived ECMs showed a more heterogeneous composition. In contrast, the dermis, epidermis and muscle ECMs were more restricted in abundant proteins. Native proportions were particularly well retained after the decellularization step of artery and epidermis. Remarkably, α1 chains of collagens type I (*COL1A1*) emerged as abundant in every tissue dECM, and became even more prominent in the ink state, along with α2 chains of collagens type 1 (*COL1A2*). As expected, elastin (*ELN*) appeared highly represented in the artery ink, corresponding to the elastic nature of the tissue. Aligned with previous results, the epidermal ink was most heterogeneous in composition. Together, tissue-specific signatures were partially retained after decellularization. Although relevant changes were observed in the abundance of ink-related matrisomal proteins as compared to native tissues, the final biomaterials represented complex mixtures with great potential impact on cell behavior.

**Figure 7:**
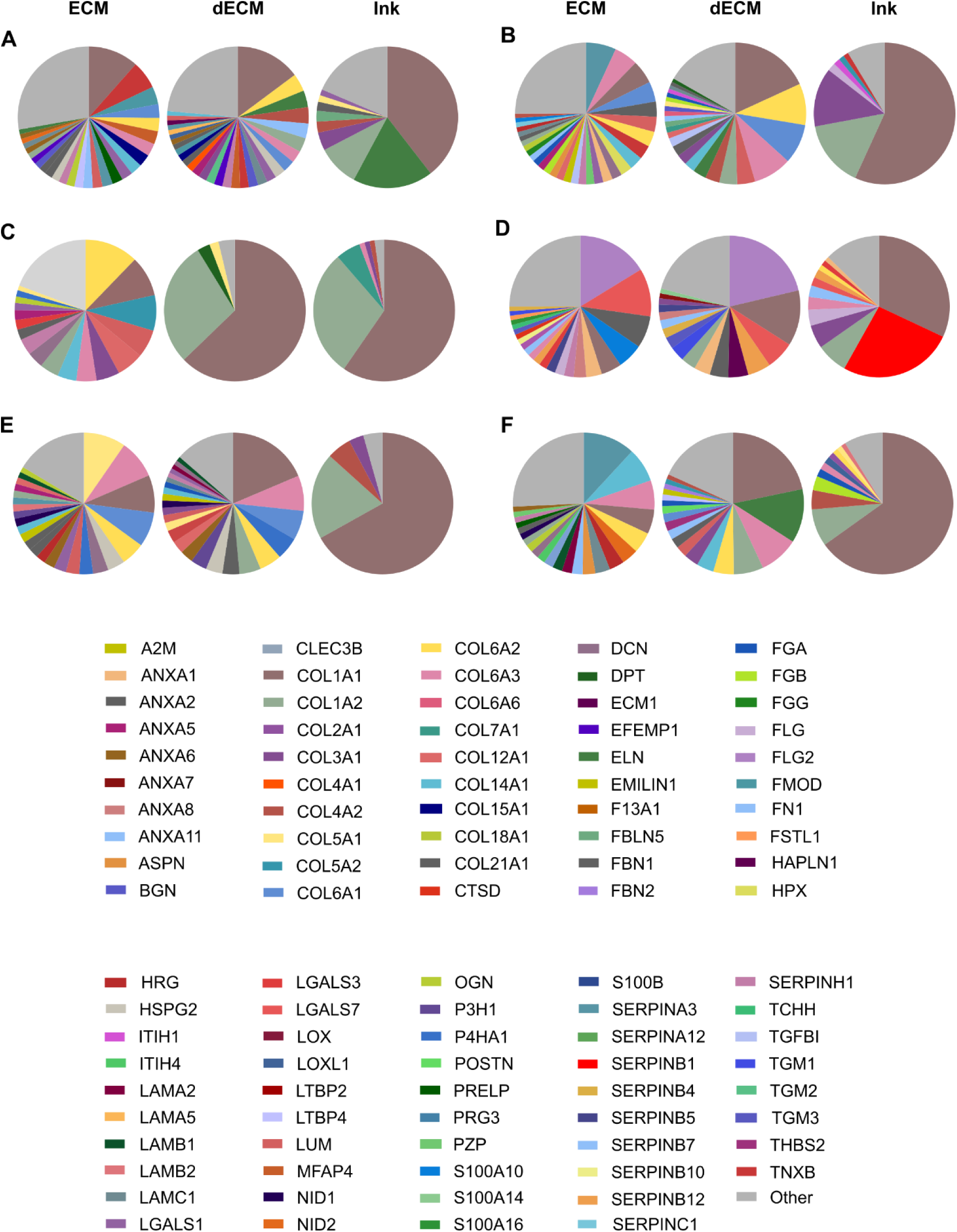
Representation of the most abundant matrisomal proteins for each tissue and sample. Identified proteins with an abundance >1% for each tissue and sample are shown for **A)** artery, **B)** breast, **C)** dermis, **D)** epidermis, **E)** muscle and **F)** nerve. Within each panel, the left, middle and right pie charts show the patterns for native tissue ECM (ECM), decellularized tissue ECM (dECM) and digested decellularized tissue ECM (Ink), respectively.

## Discussion

Due to their purported superiority in mimicking the biological and mechanical cues of the native tissues, dECM-based biomaterials are on the rise for the formulation of inks for 3D printing, as well as for direct use for regenerative medicine purposes ^11–13,43^. However, tissue decellularization processes have the potential to significantly alter sample proteomic composition, necessitating a precise evaluation of the preservation of ECM components ^44^. Notably, relatively little is known on how the different processing steps required for ink formulation affect the composition of dECM-derived inks and biomaterials ^45^. In this work, by analyzing the proteomic signature of six decellularized tissues in the three main steps of the ink formulation process (ECM, dECM, ink), we lay the foundations to better understand how ECM responds to decellularization-and digestion-associated processing, with regard to preservation of the original protein composition and ECM complexity.

The selected species was swine, for the following reasons: (i) porcine tissues are commonly available for most laboratories by sourcing them from research centers themselves and/or slaughterhouses, with limited ethical issues associated to their extraction due to their primary use for other purposes; (ii) porcine tissues share relevant anatomical and physiological similarities with human tissues, making them a relevant alternative for biomedical applications ^46,47^; (iii) these tissues offer the advantage of being a suitable source for the development of materials applicable to humans without eliciting adverse immunological responses ^48,49^. Of interest, the apparent lack of immune response seems to occur despite the presence of abundant cellular proteins in the decellularized materials ^50^. However, reports of immune response to porcine-derived dECM do exist, suggesting that elimination of some immunogenic epitopes such as α-galactosidase may suffice to avoid xenogenic rejection ^51^. In any case and as a matter of fact, most biological implants currently in use in the clinic employing dECM-derived materials are porcine-based (such as AlloDerm, Permacol and Strattice) ^47,52,53^, further demonstrating safety of human use. The selected tissues were based on easy access for tissue collection and their potential interest for purpose-built ink formulation, including major targets for tissue reconstruction efforts, such as skin, muscle and breast; and inks of use for more general purposes such as vascular ^54^ and nerve ^51^ dECM-based inks.

The characterization of matrisome components in dECM products is challenging due to their complex protein composition profiles ^55^. Traditional techniques such as immunohistochemistry, western blotting, or enzyme-linked immunosorbent assays have been applied to detect specific proteins or overall content description. However, those methods are limited to quantifying either targeted molecules or total amounts indiscriminately, and do not enable the comprehensive and high-throughput characterization of these complex materials ^23^. In this regard, LC–MS/MS technology has proven effective in deciphering the molecular fingerprint of tissues as well as the composition of the matrisome, with numerous benefits, including high sensitivity, diverse sample analysis capabilities, and low detection limits ^4^.

Regarding the ink composition, collagens and proteoglycans seemed to be the best-preserved proteins independently of the tissue, while the matrisome-associated groups were the most altered by ink production steps. In fact, it has been described that large macromolecules such as those comprising the core matrisome, have increased processing resilience because of their relatively large size and the existence of intermolecular cross-links. Smaller constituents such as growth factors, chemokines, and other signaling molecules that are newly synthesized and not yet cross-linked are largely removed ^56,57^.

The PLS-DA and VIP score analyses evidenced the variation in the protein composition of the samples according to the processing stages. In accordance with the previous results, the ECM and dECM samples tended to share more similarities due to the better preservation of the protein profile, while the inks, as a result of the decrease in the number and abundance of proteins, tended to have a relatively distinct protein profile. In addition, the intrinsic variability of biological samples and the dynamic nature of ECMs may have increased this effect. Even so, there are cases in which the proteins are relatively more abundant in decellularized samples and in the derived inks, possibly due to a masking effect of the proteins. This effect has been described on numerous occasions, where the high abundance of collagen peptides hinders the detection of other less abundant but functionally important proteins ^4^.

Additionally, fluctuations among the final materials may be shaped by factors such as the characteristics of the samples, inherent variations in the experimental procedures, and the LC-MS/MS method itself. In fact, sample digestion and extraction for MS has proven to be a challenging task due to the inherent complexity, heavy glycosylation and crosslinking, and general insolubility of ECM proteins, often leaving behind an insoluble pellet, which might result in incomplete characterization ^58,59^. Moreover, most tissues are innervated and contain blood vessels, which are not totally removed in the decellularization process. Nevertheless, it is possible to mitigate this diversity by creating pools of dECM powder for individual tissues, all the while preserving the biological significance of the materials. Recently, Biehl *et al.* developed an automated, standardized decellularization protocol that was applied to produce ECM-based hydrogels from six independent porcine tissues ^60^. Our approach fundamentally differs from theirs in that we pursued the best decellularization protocol for each tissue, thus optimizing ECM yield, but possibly at a price of increasing inter-tissue variability. Moreover, this study does not include the characterization of the final dECM-derived inks in terms of proteomic composition.

The development of biomimetic biomaterials entails an important step in basic and applied research, as well as for direct treatment applications based on tissue engineering. In this study, we presented six tissue-derived biomaterials with complex and heterogeneous biochemical compositions. Even if a proteomic loss occurred in the step from dECM to dECM-derived inks (which might be attributed to the digestion of the dECM powder), still, many proteins were identified. Interestingly, tissue-specific proteins and signatures were highlighted for every tissue. Strikingly, the proportions of the matrisomal biomolecules were certainly unique. Indeed, the relation of the presence of biomolecules within such materials with their biological function, which is commonly ignored, was addressed in this study. In all the inks, at least 45 ECM-related proteins were identified, which are closely involved in biological processes such as cell migration, cell-ECM interactions, adhesion, and proliferation. This study represents a relevant resource for investigators interested on the development of novel tissue engineering-based solutions that better mimic the native tissue environment for future clinical applications.

## Methods

### Tissue decellularization and formulation of dECM-based inks

Porcine tissue collection was sourced from cadaveric animals according to Royal Decrees 118/2021 and 53/2013 and the three Rs principles, to ensure the protection of animals used in experiments and other scientific purposes. Tissue samples were collected postmortem from 2 months old Large White female pigs sacrificed either for surgical training of clinicians at the BioGipuzkoa Health Research Institute animal facility or for food consumption at the Basatxerri slaughterhouse. Once cleaned of surrounding tissue, sample tissues were sliced and stored at -80 °C until use. Porcine breast tissue was decellularized as described previously ^27^. Dermis and epidermis were isolated from porcine dorsal skin and decellularized, as detailed in earlier research ^28^.

For aortic artery, *biceps femoris* and sciatic nerve sample decellularization, tailored protocols were developed in accordance with the recommendations given by Crapo and collaborators ^26^, as compiled in **Table S1**. Briefly, samples were freeze-thawed and exposed to a hypertonic solution based on sodium chloride, ethylenediaminetetraacetic acid (EDTA) and Tris overnight (O/N). Following, the three tissues were incubated in a detergent solution for 48 h. Specifically, artery and muscle samples were exposed to a sodium dodecyl sulfate (SDS) and sodium deoxycholate (SDC) solution, while nerve samples to a Tris/EDTA/SDS buffer. Then tissues were incubated in a nuclease (DNARASE, c-LEcta) treatment O/N and finally in a hypertonic solution based on Tris/NaCl for 4 h. A final step of sterilization by using a combination of ethanol (EtOH) and peracetic acid (PAA) for 2 h and a final 72-hour washing step with phosphate buffered saline (PBS) were performed. Samples were then stored at -80 °C. Tissue decellularization was verified by haematoxylin-eosin staining, DAPI and DNA quantification as described in ^28^, results are shown in **Figure S1**.

For obtaining the dECM inks, samples were freeze-dried (Alpha 2-4 LSCplus, Christ) and pulverized in a Pulverisette 14 premium line mill (Fritsch). The resulting powder was digested under constant agitation as described in **Table S2**. The digests were then neutralized on ice to pH 7.4 and stored at 4°C until use.

### Liquid Chromatography-Mass Spectrometry proteomics

The proteomic analyses were performed by the Proteomics Platform of CIC bioGUNE (Derio, Spain). In total, three biological replicates of ECM, dECM and inks of each tissue were analyzed. Samples were incubated in a solution containing 7 M urea, 2 M thiourea, 4 % (w/v) 3-[(3-cholamidopropyl)dimethylammonio]-1-propanesulfonate (CHAPS) and 5 mM dithiothreitol (DTT) for 30 min at RT under agitation and digested following the filter-aided sample preparation (FASP) protocol ^61^, with minor modifications. Trypsin was added to a trypsin:protein ratio of 1:50, and the mixture was incubated overnight at 37 °C, dried out in a *RVC2 25* speedvac concentrator (Christ), and resuspended in 0.1 % formic acid (FA). Peptides were desalted and resuspended in 0.1 % FA using C18 stage tips (Millipore).

Samples were analyzed in a hybrid trapped ion mobility spectrometry – quadrupole time of flight mass spectrometer (timsTOF Pro with PASEF, Bruker Daltonics) coupled online to a *nanoElute* liquid chromatograph (Bruker). 200 ng of each sample were directly loaded in a 15 cm *Bruker nanoelute FIFTEEN C18* analytical column (Bruker) and resolved at 400 nL/min with a 100 min gradient. Column was heated to 50 °C using an oven.

Protein identification and quantification was carried out using MaxQuant software using default settings. Searches were carried out against a database consisting of pig protein entries (Uniprot/Swissprot+TrEMBL/BLAST), with precursor and fragment tolerances of 20 ppm and 0.05 Da. Contaminants such as keratins and other proteins were discarded, and only proteins identified with ≥2 peptides at False Discovery Rate (FDR) <1 % were considered for further analysis.

### Data processing

Once the identifications were conducted, intracellular proteins were discarded, and only ECM-related proteins were plotted using GraphPad Prism 8.3.0 software and analyzed according to MatrisomeDB categories ^29^. The mean and standard deviation (SD) for each sample (ECM, dECM and ink) of the Label-Free Quantification (LFQ) data were calculated and normalized to the total intensity value for each sample. Venn diagrams were done using Flaski 3.12.2 ^62^and InteractiVenn ^63^. Results related to gene ontologies (GO Biological Process 2023) were obtained with Enrichr ^64^. For that, matrisome proteins of each sample were loaded and parameters were set in order to include the top 500 most relevant proteins encoding genes. The results were then sorted according to the adjusted *p*-value and the top 10 concepts were plotted in each case. PLS-DA, VIP scores and heatmap analysis were obtained with Metaboanalyst 5.0 software ^65^ by using one factor statistical analysis. Data was auto-scaled and analysed without taking into account the order of groups.Cross validations were performed with the same software in order to estimate the predictive ability of the PLS-DA models, for that 5-fold CV method was used (**Figure S6**). In cross validation analysis, the accuracy parameter represents the proportion of correctly classified observations. R^2^ indicates the explained variance, meaning the amount of variance in the data that the model can explain. Q^2^ represents the predictive ability or power of the model, and is a measure of how well the model predicts the dependent variable during cross validation. All three parameters are represented in a range of values from 0-1, with values greater than 0.5 and close to 1 having the highest predictive power. VIP scores greater than 1 were used as a threshold to identify important variables. Biological interpretation of the results was conducted using GeneCards ^66^ and The Human Protein Atlas ^67^. The workflow for the processing and analysis of the data is shown in **Figure S7**.

## Author contributions

**AIr & PV-A**: Conceptualization, Methodology, Software, Validation, Formal Analysis, Investigation, Data Curation, Visualization, Writing - Original Draft, Writing - Review & Editing. **LZO:** Data curation. **PG and KdlC**: Funding acquisition, Resources, Supervision, Project administration, Writing - Review & Editing. **AIz**: Conceptualization, Funding acquisition, Resources, Supervision, Project administration, Writing - Original Draft, Writing - Review & Editing. All authors read and approved the final manuscript.

## Acknowledgements

We thank Basatxerri for their support in sourcing fresh porcine tissue for the study. We express our gratitude to Prof. J. Ruiz Cabello and Dr. P.S. Valera for their valuable discussions regarding the analysis tools. This work was funded by Instituto de Salud Carlos III (ISCIII) and co-funded by the European Union (grants PI19/01621, PI22/01247, PT23/00142 and DTS24/00167), and Elkartek (bMG24; KK-2024/00041) and Hazitek grants (ITEAS; ZE-2022/00021) from the Department of Economic Development, Sustainability and Environment of the Basque Government. AIr was funded by a fellowship of the Predoctoral Training Program for Non-PhD Research Staff (PRE_2019_1_0031) of the Department of Education of the Basque Government and by BioGipuzkoa HRI. PV-A was supported by a fellowship paid by a donation made by Asociación Katxalin to BioGipuzkoa HRI in 2019, and by CIC biomaGUNE. The funders played no role in study design, data collection, analysis and interpretation of data, or the writing of this manuscript.

## Data availability

The mass spectrometry proteomics data have been deposited to the ProteomeXchange Consortium via the PRIDE partner repository with the dataset identifier PXD059171. The datasets analyzed during the current study are available in Zenodo (10.5281/zenodo.14195914). The workflow and metadata are available at protocols.io (dx.doi.org/10.17504/protocols.io.8epv52zr5v1b/v1).

