## Supplementary Data for "Comparative proteomic analysis of the composition of decellularized extracellular matrix (dECM) and dECM-based inks as compared to the native tissue"

**Supplementary Figures**

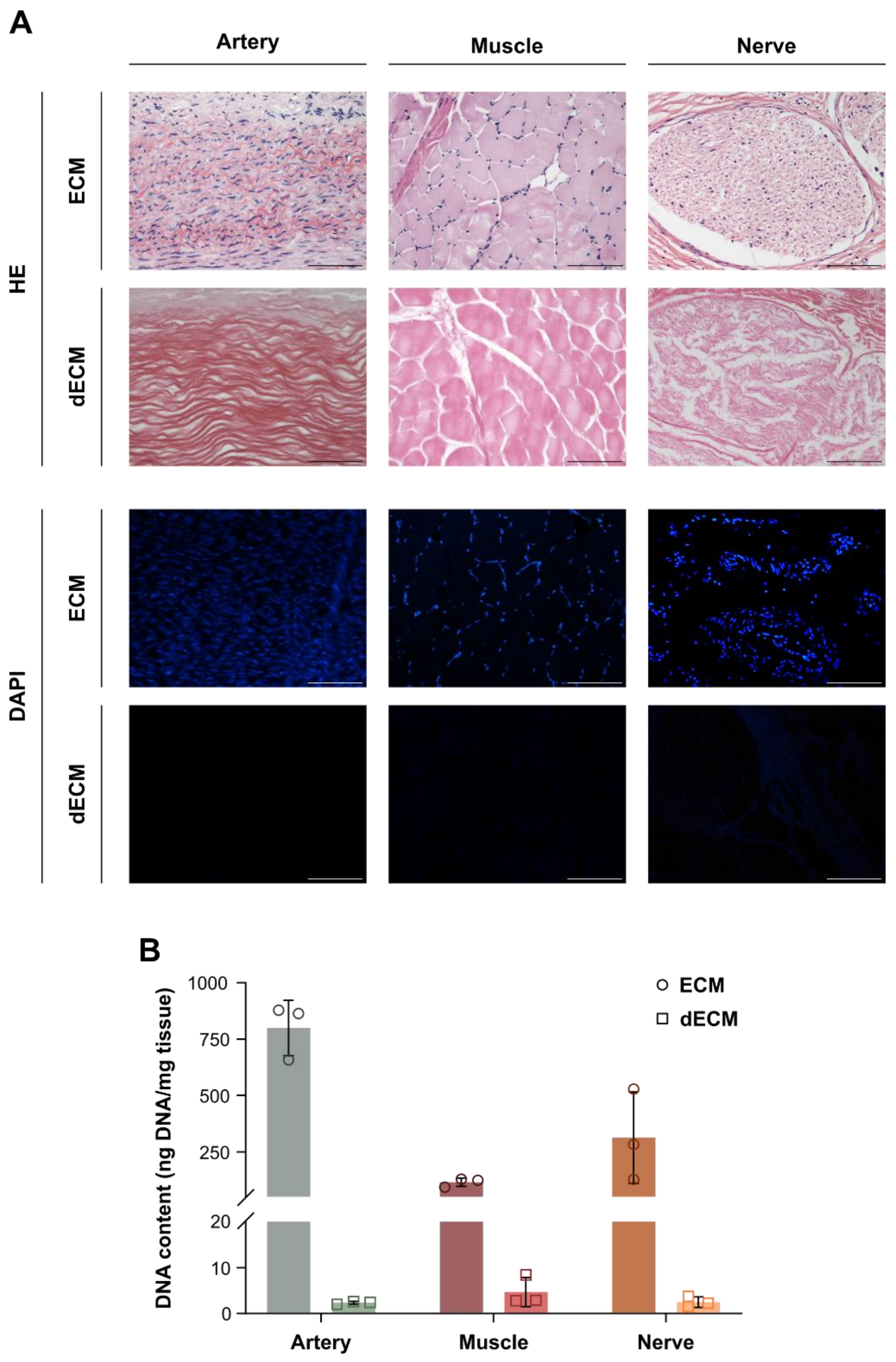


***Figure S1.*** *Characterization of tissue and decellularized extracellular matrices (dECM) derived from decellularization of artery, muscle and nerve tissues.* ***A)*** *Histological analysis of extracellular matrices (ECMs) and dECMs, comprising haematoxylin and eosin and DAPI fluorescence staining. Scale bars: 100 µm. B) Quantification of the DNA content of native and decellularized ECMs.*

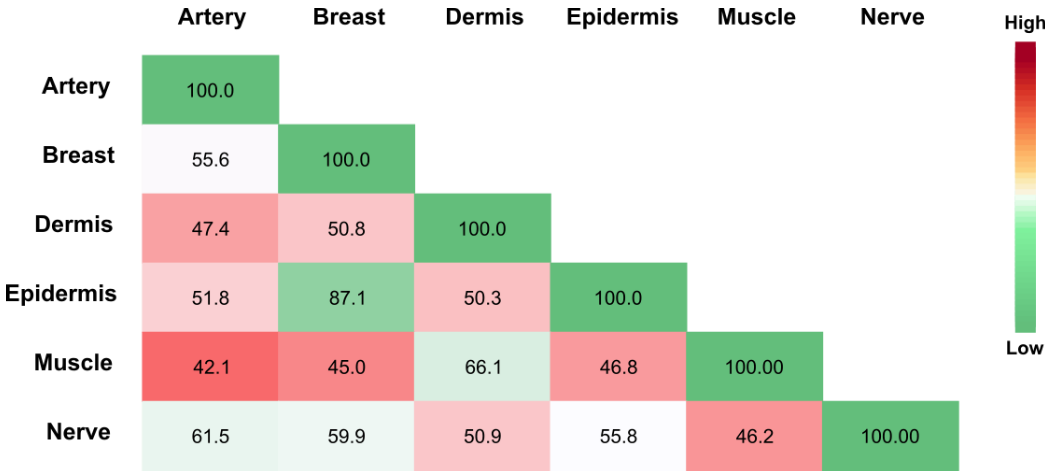


***Figure S2.*** *Heatmap of the matrisome proteins coverage shared between native tissues.*

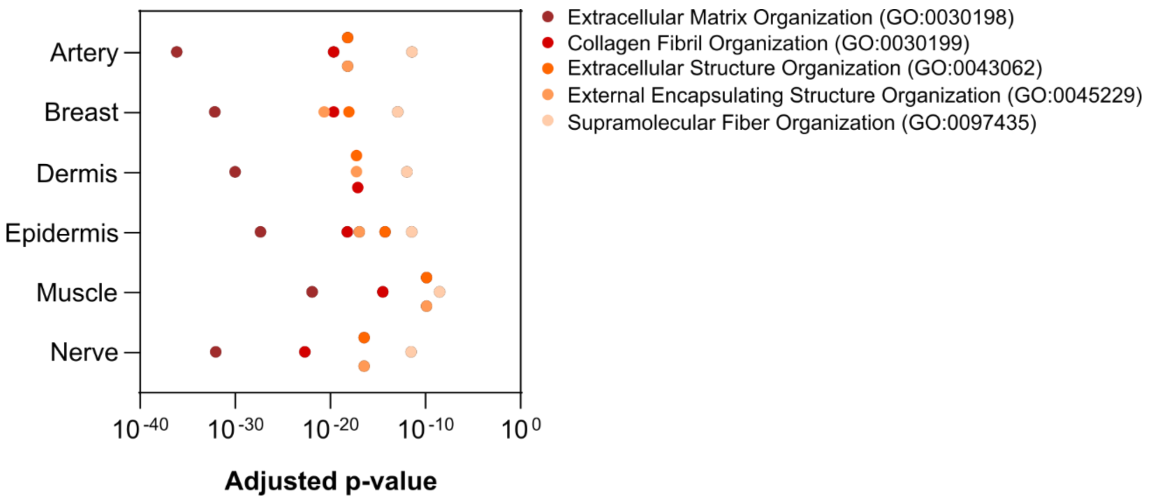


***Figure S3.*** *Adjusted p-values of biological processes gene ontology (GO) common to every tissue ECM.*

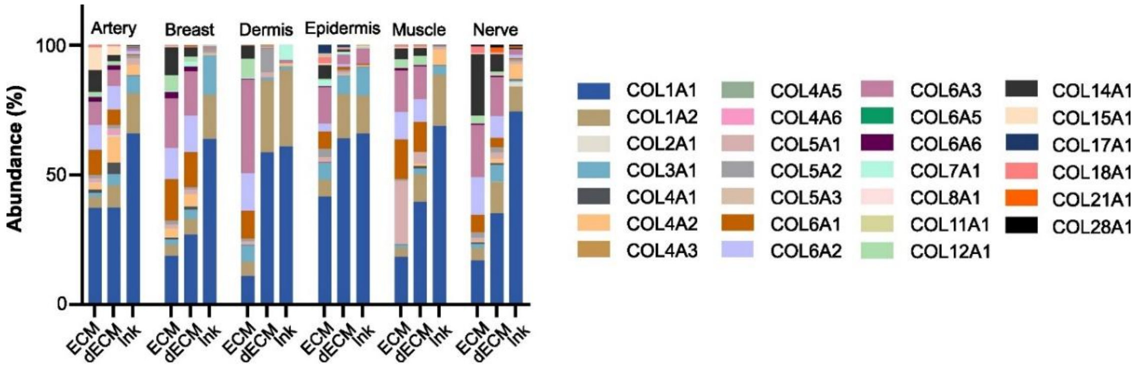


***Figure S4.*** *Collagen types present within the collagen family for each tissue ECM, dECM and ink.*

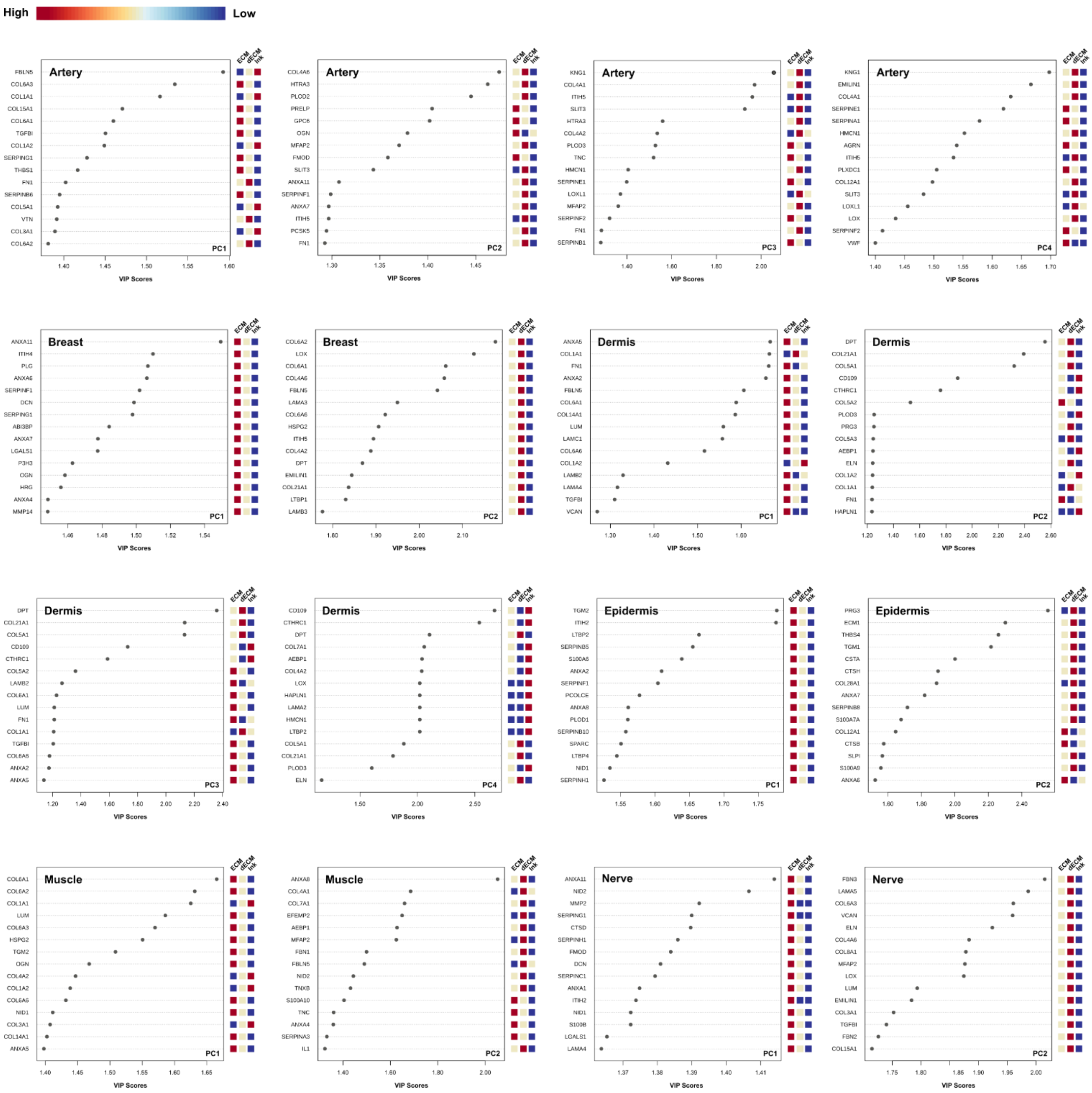


***Figure S5.*** *Partial Least-Squares Discriminant Analysis (PLSDA)-based Variable Influence/Importance on Projection (VIP) scores of artery, breast, dermis, epidermis, muscle and nerve ECM, dECM and Ink for each representative principal component (PC).*

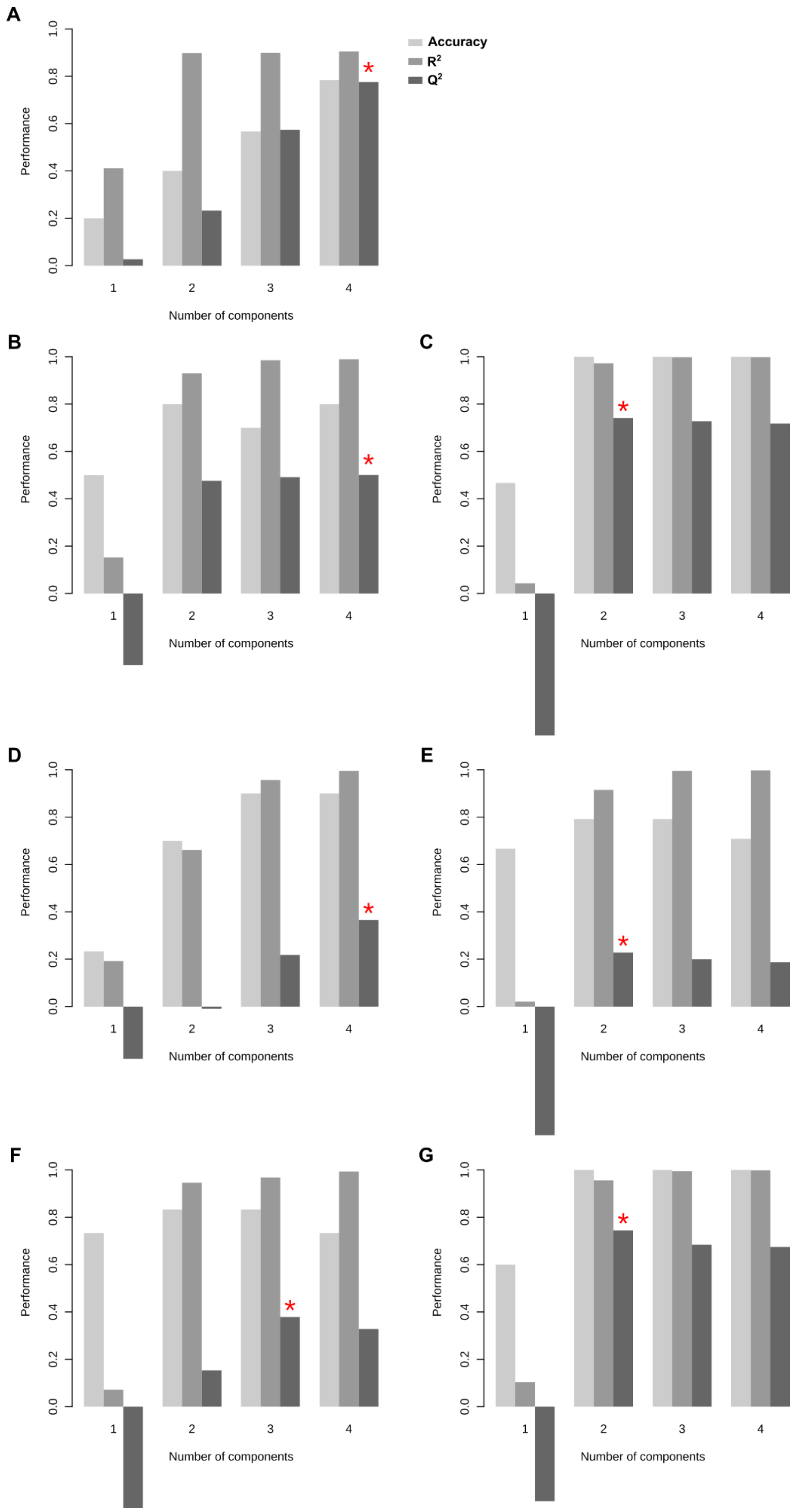


***Figure S6.*** *Cross validation of the PLS-DA results of* ***A)*** *ECMs,* ***B)*** *artery,* ***C)*** *breast,* ***D)*** *dermis,* ***E)*** *epidermis,* ***F)*** *muscle and* ***G)*** *nerve. Accuracy (*) represents the best model based on Q2 results.*

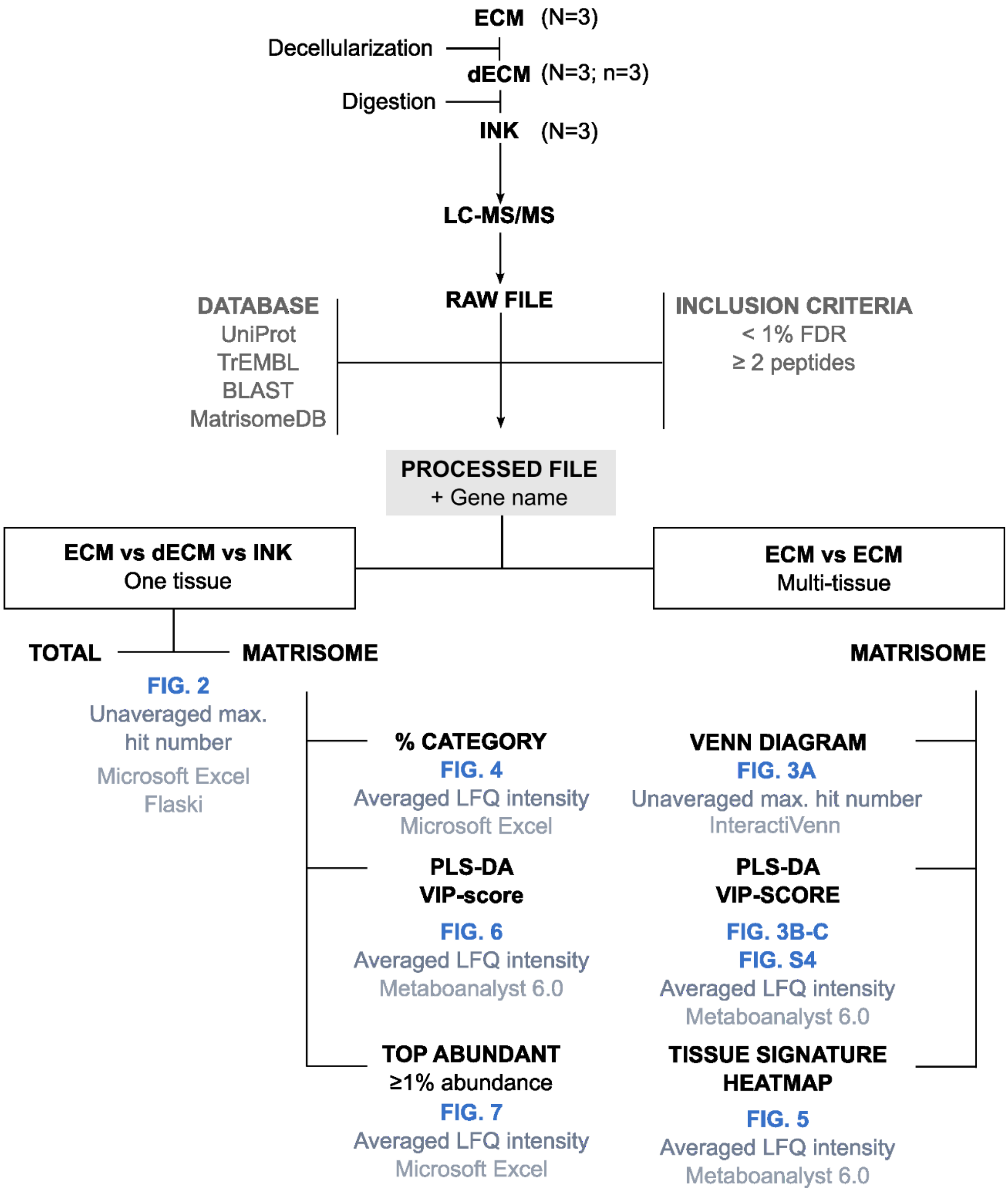


***Figure S7.*** *Workflow for the processing and analysis of the data.*

**Supplementary Tables**

***Table S1.*** *Decellularization protocol for aortic artery (A), biceps femoris muscle (M) and sciatic nerve (N).*

| Solution | Time | Temperature | Agitation |
|---|---|---|---|
| 10 mM Tris + 1% EDTA + 1.5 M NaCl | ON | RT | 180 rpm |
| **(A)** 0.5% SDS + 0.5% SDC<br>**(M)** 1% SDC + 1% SDS<br>**(N)** 10 mM Tris + 0.1% EDTA + 1% SDS | 48 h | RT | 180 rpm |
| 50 mM Tris + 1.5 M NaCl | 6 h | RT | 180 rpm |
| 40 U/ml DNARASE Buffer | ON | 37ºC | 180 rpm |
| 50 mM Tris + 1.5 M NaCl | 4 h | RT | 180 rpm |
| 0.1% PAA + 4% EtOH | 2 h | RT | 60-100 rpm |
| PBS 1x | 72 h | 4ºC | 60-100 rpm |

** EDTA,* ethylenediaminetetraacetic acid*; SDS, sodium dodecyl sulfate, SDC, sodium deoxycholate; PAA, peracetic acid; EtOH, ethanol; PBS, phosphate buffered saline solution; O/N, overnight; h, hours; RT, room temperature; rpm, revolutions per minute.*

***Table S2.*** *Digestion conditions for the different tissues.*

| Tissue | Concentration (mg/mL) | Solvent | Pepsin (1 mg/mL) | Time (h) |
|---|---|---|---|---|
| Artery | 10 | 0.5 M acetic acid | Yes | 48 |
| Nerve | 10 | 0.5 M acetic acid | Yes | 48 |
| Muscle | 10 | 0.5 M acetic acid | Yes | 48 |
| Dermis | 10 | 0.5 M acetic acid | Yes | 72 |
| Epidermis | 10 | 10 M sodium hydroxide | No | 48 |
| Breast | 25 | 0.5 M acetic acid | Yes | 48 |

***Table S3.*** *Quantification of the number and proportion of the total as well as matrisome proteins for each tissue and sample.*

| | TOTAL PROTEINS [N (% of ECM)] | | | MATRISOMAL PROTEINS [N (% of ECM)] | | |
|---|---|---|---|---|---|---|
| **SAMPLE** | **ECM** | **dECM** | **Ink** | **ECM** | **dECM** | **Ink** |
| **Artery** | 2529 (100) | 2610 (103.2) | 477 (18.9) | 170 (100) | 172 (101.2) | 61 (35.9) |
| **Breast** | 2483 (100) | 1819 (73.3) | 672 (27.1) | 179 (100) | 164 (91.6) | 94 (52.5) |
| **Dermis** | 1529 (100) | 666 (43.6) | 321 (21.0) | 109 (100) | 58 (53.2) | 45 (41.3) |
| **Epidermis** | 2513 (100) | 2557 (101.8) | 732 (29.1) | 169 (100) | 176 (104.1) | 122 (72.2) |
| **Muscle** | 1209 (100) | 1228 (101.6) | 321 (26.6) | 83 (100) | 85 (102.4) | 39 (47.0) |
| **Nerve** | 2131 (100) | 1495 (70.2) | 203 (9.5) | 152 (100) | 128 (84.2) | 45 (29.6) |
| **Median** | 2307 (100) | 1657 (87.0) | 399 (23.8) | 160 (100) | 146 (96.4) | 53 (44.1) |
| **Range** | 1209-2529 (100-100) | 666-2610 (43.6-103.2) | 203-732 (9.5-29.1) | 83-179 (100-100) | 58-176 (53.2-104.1) | 39-122 (29.6-72.2) |
| **SD** | 568.4 (0) | 762.3 (24.2) | 211.5 (7.3) | 38.8 (0) | 49.5 (19.3) | 33.3 (15.0) |

** N, number; SD, standard deviation.*

***Table S4**. GO terms, adjusted p-values and related gene names for each tissue ECM.*

| | GO term | Adjusted p-value | Genes |
|---|---|---|---|
| **Artery** | Regulation Of Plasminogen Activation (GO:0010755) | 7,46E-07 | F12; CTSZ; SERPINE1; THBS1 |
| | *Positive Regulation Of Wound Healing (GO:0090303)* | 1,79E-05 | SMOC2; F12; SERPINE1; THBS1 |
| | Positive Regulation Of Hemostasis (GO:1900048) | 1,79E-05 | F12; SERPINE1; THBS1 |
| **Breast** | Fibrinolysis (GO:0042730) | 1,09E-05 | FGA; SERPINF2; F2 |
| | Negative Regulation Of Blood Coagulation (GO:0030195) | 1,00E-04 | FGA; SERPINF2; F2 |
| | Negative Regulation Of Fibrinolysis (GO:0051918) | 1,66E-03 | SERPINF2; F2 |
| | Positive Regulation Of Collagen Biosynthetic Process (GO:0032967) | 1,80E-03 | SERPINF2; F2 |
| **Epidermis** | Negative Regulation Of Endoeptidase Activity (GO:0010466) | 1,44E-10 | SERPINB10; SERPINB13; SERPINB2; SERPINB7; SERPINB8; SERPINB5; ANXA8 |
| | Negative Regulation Of Peptidase Activity (GO:0010951) | 5,47E-09 | SERPINB10; SERPINB13; SERPINB2; SERPINB7; SERPINB8; SERPINB5 |
| | Regulation Of Peptidase Activity (GO:0010951) | 8,67E-09 | SERPINB10; SERPINB13; SERPINB2; SERPINB7; SERPINB8; SERPINB5 |
| | Epidermis Development (GO:0008544) | 1,26E-04 | COL17A1; FLG2; LAMA3; TGM5 |
| **Muscle** | Regulation Of Basement Membrane Organization (GO:0110011) | 2,25E-14 | LAMA2; LAMB1; LAMB2; NID1 |
| | Positive Regulation Of Integrin-Mediated Signaling Pathway (GO:2001046) | 1,11E-12 | LAMA2; LAMB1; LAMB2; NID1 |
| | Positive Regulation Of Muscle Cell Differentiation (GO:0051149) | 2,05E-12 | LAMA2; LAMB1; LAMB2; NID1 |
| | Regulation Of Integrin-Mediated Signaling Pathway (GO:2001044) | 4,36E-12 | LAMA2; LAMB1; LAMB2; NID1 |
| **Nerve** | Extracellular Matrix Organization (GO:0030198) | 8,89E-06 | COL28A1; COL14A1; ADAM10; NID2; MATN2 |
| | Extracellular Structure Organization (GO:0043062) | 1,79E-03 | COL28A1; COL14A1; MATN2 |
| | *External Encapsulating Structure Organization (GO:0045229)* | 1,79E-03 | COL28A1; COL14A1; MATN2 |
| | Wnt Signaling Pathway (GO:0016055) | 2,99E-03 | WNT5B; GPC4 |
